# Transfer Learning Enables Drug–Target Interaction Prediction in Data-Scarce One-Carbon Metabolism

**DOI:** 10.64898/2026.04.30.721937

**Authors:** Alperen Dalkiran, Takugo Cho, M. Volkan Atalay, Kun Woo D. Shin, Angelo Y. Meliton, Yufeng Tian, Parker S. Woods, Obada R. Shamaa, Robert B. Hamanaka, Gökhan M. Mutlu, Rengul Cetin-Atalay

## Abstract

Predicting drug–target interactions (DTIs) with deep learning offers opportunities to accelerate drug discovery, yet performance is constrained by the scarcity of target-specific training data. This is a particular challenge for mitochondrial one-carbon (1C) pathway enzymes, which are attractive therapeutic targets but remain pharmacologically understudied. Mitochondrial 1C metabolism supplies glycine, reducing equivalents, and 1C units critical for nucleotide synthesis, and has emerged as a key pathway in cancer and fibrosis. SHMT2 and MTHFD2, two key 1C enzymes, support collagen production in fibroblasts, blocking either prevents TGF-β–induced glycine and collagen accumulation. Here, we developed transfer learning–based deep learning models to predict interactions between approved drugs and SHMT2 or MTHFD2 despite minimal target-specific training data, pre-training on large datasets from related enzymes before fine-tuning to these targets. Virtual screening of the DrugBank library identified six candidates, three of which, Carbimazole, Crizotinib, and GSK2018682 reduced TGF-β–induced collagen production and glycine accumulation in human lung fibroblasts, demonstrating transfer learning as a strategy for repurposable drug identification in data-scarce metabolic targets.

## INTRODUCTION

In the evolving field of pharmacology, predicting drug-target interactions (DTIs) is pivotal for identifying therapeutic candidates efficiently and cost-effectively. Traditionally, drug discovery is a long, laborious and expensive process (Vamathevan *et al*, 2019). As an alternative, artificial intelligence-powered computational approaches recently emerged, offering the potential to significantly accelerate this process. A unique challenge of machine learning-based DTI prediction is its need for specificity (Rifaioglu *et al*, 2019). Each target protein or protein family interacts uniquely with potential drug molecule ligands, requiring a customized computational model for each. To address this, Deep Neural Network (DNN) models have been developed for each target protein or family (Iskar *et al*, 2013; Rifaioglu *et al*, 2020; Trepte *et al*, 2024). These models are specifically engineered to assess the biological activity of compounds, effectively classifying them based on their interaction potential with designated targets.

Drug-target interaction prediction is a ligand-based binary classification problem, where deep neural networks use drug compound features to categorize ligands as either active or inactive against a specific protein(Rifaioglu *et al*., 2020; Vamathevan *et al*., 2019). The challenge of classification is often addressed by developing a unique deep neural network model for each target protein or protein family. The bioactivity prediction accuracy of deep learning models depends on the size of the training data, which is often unavailable for many target proteins or protein families. Therefore, predictive models are lacking for most proteins. Given the limitation of data availability in the domain of specific protein-ligand interactions, there is a need for DNN approaches which incorporates advanced machine learning strategies such as transfer learning (Gangwal *et al*, 2024). Transfer learning enables the use of models pre-trained on extensive, general datasets for new, specific tasks allowing the adapted models to operate with improved accuracy and efficiency, even when only limited data specific to the task is available.(Dalkıran *et al*, 2023; Zhao *et al*, 2025)

The mitochondrial one-carbon (1C) pathway has recently received increased attention as a target for drug discovery (Dekhne *et al*, 2020; Liu *et al*, 2025). This pathway is an important cellular source of glycine, reducing equivalents, and 1C units which can be exported from the mitochondria and used to produce nucleotides in the cytoplasm (Ducker & Rabinowitz, 2017; Locasale, 2013; Tibbetts & Appling, 2010). The first two enzymes of the mitochondrial 1C pathway, serine hydroxymethyltransferase 2 (SHMT2) and 5,10-methylene tetrahydrofolate dehydrogenase 2 (MTHFD2) are independent prognostic indicators for multiple cancers (Kim *et al*, 2015; Liu *et al*, 2014; Liu *et al*, 2019; Noguchi *et al*, 2018; Tibbetts & Appling, 2010). SHMT2 and MTHFD2 have also been linked to development of fibrotic disease as fibroblasts upregulate mitochondrial 1C metabolism to support the synthesis of glycine, the most abundant amino acid in collagen matrix proteins (Meliton *et al*, 2025a). We have previously demonstrated that inhibition of either SHMT2 or MTHFD2 prevents increases in cellular glycine concentration after stimulation of lung fibroblasts with the profibrotic cytokine transforming growth factor-β (TGF-β). Thus, the development of therapeutics targeting mitochondrial 1C metabolism may be valuable in multiple disease settings.

Here we demonstrate the use of deep transfer learning to predict drug-target interactions (DTIs) for understudied targets within one carbon pathway tandem enzymes SHMT2 and MTHFD2, despite having scarce training data. Given the limited data availability for our specific task, we employed transfer learning techniques, leveraging the adaptability of deep transfer learning. This study presents the effectiveness of transfer learning by training a neural network classifier on a large, generalized source dataset and then fine-tuning it on smaller, specialized targets: SHMT2 and MTHFD2. Our results show that transfer learning significantly identifies drugs that act on one carbon pathway enzymes, which are validated *in vitro*. Additionally, we demonstrated the value of the transfer learning method and highlighted its potential for DTI prediction in scenarios with limited data, such as one-carbon pathway tandem enzymes MTHFD2 and SHMT2.

## RESULTS

### MTHFD2 and SHMT2 related DTI data construction for transfer learning model

The datasets for targeting the critical enzymes MTHFD2 and SHMT2 of mitochondrial 1C pathway, were constructed from the ChEMBL bioactive molecule repository (Gaulton et al. 2012). Our approach included the rigorous compilation of positive and negative training and test data sets for both our targets (SHMT2 and MTHFD2) and the other enzymes within the specific NC-IUBMB-Enzyme Classification (EC) nomenclature class. The EC nomenclature class in the tree structure is used to construct the training and test datasets for source and target domains (Table 1 and Table 2). As a bifunctional methylenetetrahydrofolate dehydrogenase/cyclohydrolase, MTHFD2 is associated with two EC numbers EC:1.5.1.15, NAD-dependent methylenetetrahydrofolate dehydrogenase and EC:3.5.4.9, methenyltetrahydrofolate cyclohydrolase. By applying strict filtering criteria based on target specificity, assay reliability, and chemical diversity, we aimed to reduce bias and ensure robust MTHFD2 DNN model training. The *target positive dataset* for the MTHFD2 then contained 29 unique compounds which is a number very low to train traditional DNN from scratch (Figure 1B). IUBMB-EC Transferases (EC:2.–.–.–) class enzymes and their interacting compounds were used to form inactive compound set for MTHFD2 and 29 compounds were randomly selected from this set as the *target (protein) negative dataset*. The positive and negative datasets were balanced. 50 training sets were formed by randomly selecting 25 compounds from each target protein dataset (to create *target training dataset*), with the remaining 8 compounds assigned to the *target test dataset*.

**Table 1.** MTHFD2- model test and training data sets.

|  | *Name* | *# of DTI* | *# of unique DTI* | *active* | *clustered* |
| --- | --- | --- | --- | --- | --- |
| **MTHFD2** | Bifunctional methylenetetrahydrofolate dehydrogenase / cyclohydrolase, mitochondrial | 40 | 40 | 40 | 29 |
| 1.5.1.15 | NAD-dependent methylenetetrahydrofolate dehydrogenase | - | - | - | - |
| 1.5.1.- | With NAD or NADP as acceptor | 3 368 | 2 223 | 822 | 578 |
| 1.5.-.- | Acting on the CH-NH group of donors | 3 385 | 2 240 | 827 | 583 |
| 1.-.-.- | Oxidoreductases | 60 368 | 46 190 | 14 909 | 10 280 |
| 3.5.4.9 | Methenyltetrahydrofolate cyclohydrolase | - | - | - | - |
| 3.5.4.- | Cyclic Amidines | 557 | 455 | 139 | 99 |
| 3.5.-.- | Acting on carbon-nitrogen bonds, other than peptide bonds | 4 110 | 3 265 | 1 204 | 722 |
| 3.-.-.- | Hydrolase Nomenclature | 183 413 | 154 863 | 54 293 | 30 311 |

**Table 2.** SHMT2- model test and training data sets.

|  | *Name* | *# of DTI* | *# of unique DTI* | *active* | *clustered* |
| --- | --- | --- | --- | --- | --- |
| **SHMT2** | Serine hydroxymethyltransferase, mitochondrial | - | - | - | - |
| 2.1.2.1 | Glycine hydroxymethyltransferase | 198 | 196 | 6 | 6 |
| 2.1.2.- | Hydroxymethyl-, Formyl- and Related Transferases | 198 | 196 | 6 | 6 |
| 2.1.-.- | Transferring one-carbon groups | 3 869 | 2 117 | 1362 | 732 |
| 2.-.-.- | Transferase | 72 572 | 44 403 | 31 644 | 18 572 |

**Figure 1.**
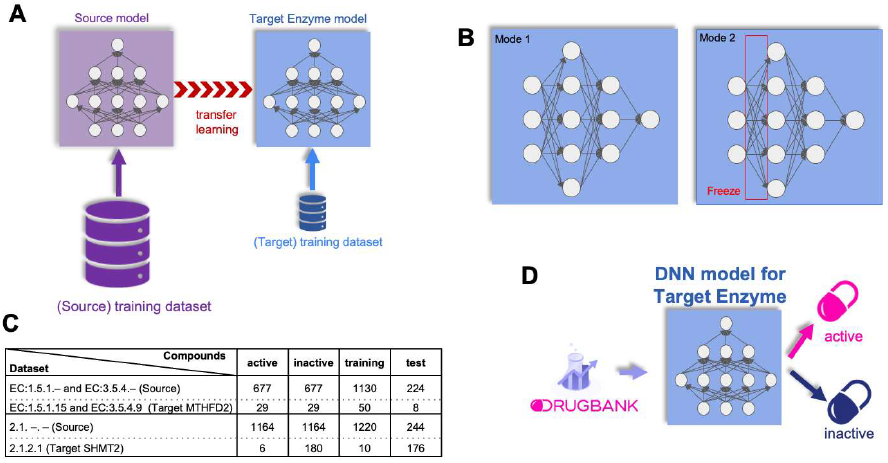
Predicting drug–one-carbon pathway enzyme interactions using transfer learning. (**A**) A source deep neural network model was first trained on a large dataset derived from a broader enzyme class (source family). The pre-trained model was then adapted to a specific target enzyme via transfer learning, using a smaller target-specific training dataset. (**B**) Two transfer learning modes were applied to a feedforward neural network (FNN): Mode 1 (full fine-tuning), in which all network weights are updated during retraining, and Mode 2 (feature transformer), in which early layers are frozen and only the later layers are retrained on the target dataset.(**C**) Composition of source and target training and test sets used for MTHFD2 and SHMT2 models. Source datasets were drawn from broader enzyme classes (EC:1.5.1.– / EC:3.5.4.– for MTHFD2; EC:2.1.– for SHMT2), while target datasets comprised compounds with confirmed activity against MTHFD2 (EC:1.5.1.15 / EC:3.5.4.9) or SHMT2 (EC:2.1.2.1). (**D**) Virtual screening of approved DrugBank compounds using the trained MTHFD2 and SHMT2 deep neural network (DNN) models, classifying each compound as active or inactive against the target enzyme.

While MTHFD2 had 29 DTI pairs available at ChEMBL repository, SHMT2 had no DTI data. Since SHMT2 and SHMT1 possess the same EC nomenclature class 2.1.2.1 and SHMT1 had 6 documented DTI data at ChEMBL repository, we used those 6 DTI data for SHMT2 DTI prediction model training. The bioactive compounds interaction with SHMT1 and MTHFD2 was selected by a pChEMBL threshold value which was set to ≤4.3 (≤50 μM) for MTHFD2 and ≤6.0 (≤1 μM for SHMT1. The *target positive dataset* for SHMT2 contained only 6 unique compounds which is too few to train a traditional DNN from scratch (Figure 1C). We therefore leveraged transfer learning, dividing *the target positive dataset* into a training set (5 compounds) and a test set (1 compound). IUBMB-EC Transferases (EC:2.–.–.–) class enzymes and their interacting compounds were used to form inactive compound set for SHMT2 (180 compounds) and 5 compounds were randomly selected from this set to form the *target negative training dataset*. The target positive and negative datasets were balanced by randomly selecting 5 compounds from each *target dataset* forming 10 *target training datasets*, with the remaining 176 (1+175) compounds assigned to the *target test dataset*.

### MTHFD2 specific and SHMT2 specific DNN transfer learning model development

The development of a MTHFD2-specific deep neural network (DNN) transfer learning model process includes two phases. In the first phase, the source neural network is trained on a dataset comprising compounds targeting EC:1.5.1.– and EC:3.5.4.– enzymes (Table 1 and Figure 1C). The pre-trained model is fine-tuned using a smaller dataset of compounds specifically targeting MTHFD2, resulting in the final MTHFD2-targeted DNN model (Figure 1).

The MTHFD2 transfer learned model resulted in an MCC value of 0.902 (GitHub page) for the source family test set, which is the performance of a deep neural network model that we trained with 1,130 compounds to predict whether a given compound interacts with the proteins in EC Numbers 1.5.1.- and 3.5.4.-that we selected as the source family. Thus, we obtained a high-quality source deep neural network model suitable for learning by transfer. For MTHFD2, if we train the model from scratch, i.e., using only 50 (25 positive and 25 negative) samples (compounds) for training, we obtain an MCC value of 0.775. However, the performance results were better (than 0.775) when we used transfer learning for MTHFD2; the MCC values were 1.0 and 1.0 for Mode 1 and Mode 2, respectively. Among the 50 models constructed for MTHFD2, several achieved MCC values of 1.0. Following a thorough assessment, we identified one model as the most suitable option, which we then employed for further predictions.

Development of serine hydroxymethyltransferase (SHMT1 or 2) specific DNN transfer learning model uses the dataset formed with the compounds acting on EC:2.1. –.– target enzymes as the *source training dataset* comprising 732 active (Table 2) and 732 inactive compounds randomly selected from EC:2.1. –.– enzymes. This data is used to pre-train a source model as a foundation for transfer learning. The pre-trained source model is then re-trained on a smaller dataset from the compounds acting on SHMT1 for fine-tuning and produce the final serine hydroxymethyltransferase target DNN model (Figure 1A and B).

We obtained an MCC value of 0.713 for the source family (test set), which is the performance of a deep neural network model that we trained with 1,464 compounds to predict whether a given compound interacts with the proteins in EC:2.1. –.– that we selected as the source family. Thus, we obtained a high-quality source deep neural network model suitable for learning by transfer. For SHMT1 or 2, if we train the model from scratch, i.e., using only 10 (5 positive and 5 negative) samples (compounds) for training, we obtain an MCC value of 0.0. When we used transfer learning for SHMT1 or 2, the MCC values were 0.2 and 0.308 for Mode 1 and Mode 2, respectively. Among the 50 models constructed for SHMT1 or 2, several achieved MCC values of 0.308. Following a thorough assessment, we identified one model as the most suitable option, which we then employed for further predictions.

### Virtual screening of DrugBank compounds using SHMT1 or 2 and MTHFD2 DNN models

Using the best model based on test set prediction performances, we screened the approved drugs of DrugBank 2 562 drugs, Version 5.1.9, 2022-01-03). The top scoring 6 drugs were Carbimazole, Leucovorin, Crizotinib, Flavoxate, Trametinib, GSK-2018682 (Figure 2 A, and full list of the DrugBank screening results is given as supplementary data). Drugs were selected based on their prediction scores, which is a probability score between 0 and 1 indicating the likelihood that the instance belongs to the positive class, on trained models of Mode 1 and Mode2 with either layer 1 or layer 2 were frozen during training. The reported mode of action of each drug varies ranging from enzyme inhibition to DNA interaction and muscle relaxation. Carbimazole is an antithyroid medication used primarily to treat hyperthyroidism. It works by inhibiting the enzyme thyroid peroxidase and reduces the production of thyroid hormones. Leucovorin (Folinic acid) is used to reduce the toxic effects of folic acid antagonists and restore normal folate function. Crizotinib is multi-kinase inhibitor reported to act on ALK and ROS1 kinases. Flavoxate is classified as a smooth muscle relaxant with mild anti-inflammatory activities. Trametinib is a kinase inhibitor acting on MAPK/ERK pathway. GSK-2018682 is an investigational small-molecule developed for the treatment of multiple sclerosis. Small molecule drugs act on signaling pathways that determine their effects, and since cell signaling involves crosstalks, these drugs may have additional bioactivities beyond their reported ones. Therefore, we tested their bioactivities *in vivo* on Serine Glycine one carbon pathway with respect to collogen synthesis.

**Figure 2.**
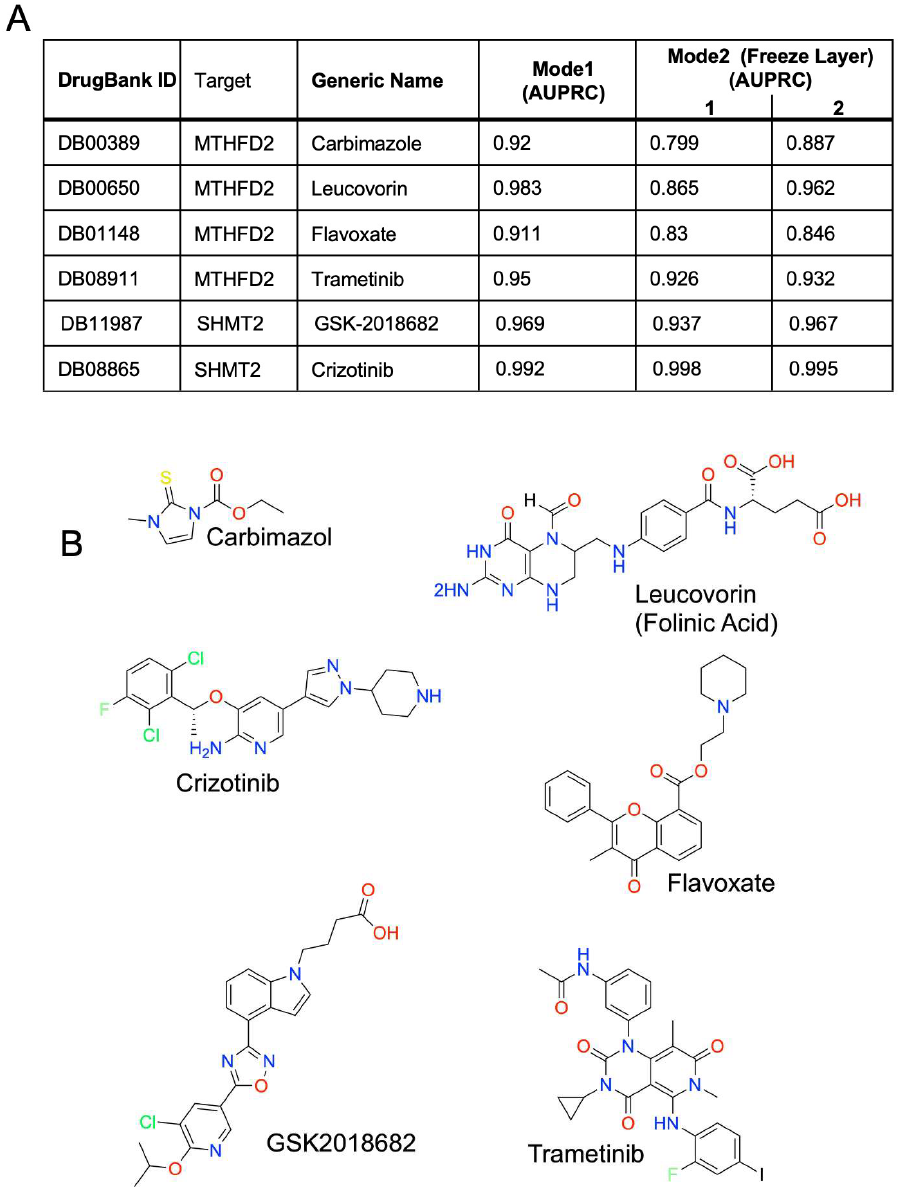
Virtual screening of DrugBank compounds identifies six candidate inhibitors of MTHFD2 and SHMT2. (**A**) Performance metrics, measured as the Area Under the Precision-Recall Curve (AUPRC), for the six top-ranked approved drugs predicted to interact with MTHFD2 (Carbimazole, Leucovorin, Flavoxate, and Trametinib) or SHMT2 (GSK-2018682 and Crizotinib), evaluated across three transfer learning configurations: Mode 1 (full fine-tuning) and Mode 2 with freeze layers 1 and 2 (feature transformer). (**B**) Chemical structures of the six candidate compounds identified by virtual screening.

We have previously shown that both SHMT2 and MTHFD2 activity are required for glycine accumulation in human lung fibroblasts after treatment with TGF-β (Meliton *et al*., 2025a; Nigdelioglu *et al*, 2016). This accumulation of glycine supports the production of glycine-rich collagen proteins which are also induced by TGF-β (Hamanaka & Mutlu, 2021a, b). We thus treated normal human lung fibroblasts (NHLFs) with TGF-β in the presence or absence of the candidate drugs. We found that Leucovorin, Flavoxate, and Trametinib did not affect collagen protein induction downstream of TGF-β; however, Carbimazole, Crizotinib, and GSK-2018682 showed dose-dependent inhibition of TGF-β-induced collagen production (Figure3A). We thus measured TGF-β-induced glycine accumulation in NHLFs treated with Carbimazole, Crizotinib, and GSK-2018682. We found that all three drugs inhibited TGF-β-induced glycine accumulation to a similar extent as the MTHFD2 inhibitor DS18561882 ((Meliton *et al*, 2025b). Together, these findings suggest that these drugs may have an inhibitory effect on the mitochondrial 1C pathway.

**Figure 3.**
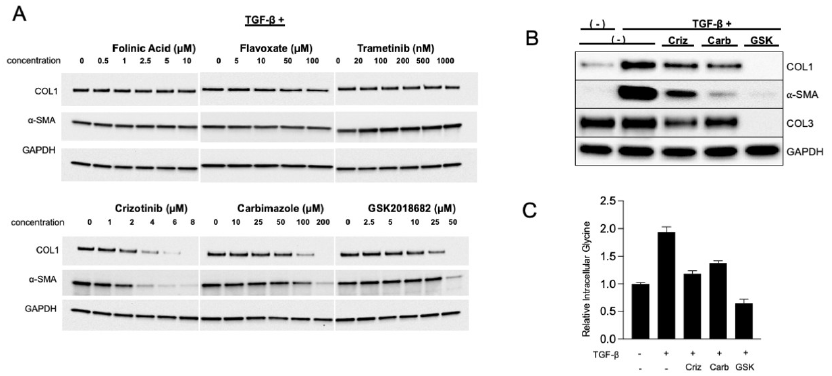
Identified drugs reduce TGF-β–induced collagen production and glycine accumulation. (**A**) Western blot analysis of collagen 1 (COL1) and α-smooth muscle actin (α-SMA) protein expression in human lung fibroblasts (HLFs) stimulated with TGF-β and treated with increasing concentrations of Folinic Acid (0–10 µM), Flavoxate (0–400 µM), Trametinib (0–1000 nM), Crizotinib (0–8 µM), Carbimazole (0– 200 µM), or GSK-2018682 (0–50 µM). GAPDH serves as a loading control. (**B**) Western blot analysis of collagen 1, collagen 3A, and α-smooth muscle actin protein expression in HLFs treated with TGF-β for 48 hours or left untreated. Cells were treated with crizotinib, carbimazole, or GSK-2018682 as indicated. (**C**) Analysis of intracellular glycine levels in HLFs treated with TGF-β for 48 hours or left untreated. Cells were treated with crizotinib, carbimazole, or GSK-2018682 as indicated.

## Discussion

For protein families, when the training dataset size is less than 100 samples, the use of transfer learning yields better results compared to starting the training process from scratch(Dalkiran *et al*., 2023). If the dataset used for training the target exceeds 100 samples, the effectiveness of transfer learning is almost equivalent to training a network model from scratch. However, transfer learning remains the preferred method as it converges with fewer training epochs. For every instance where the training dataset contained fewer than 100 samples, transfer learning outperformed the reference model. Additionally, models trained using transfer learning begin with lower initial loss values relative to the reference model. As a result, fewer epochs are typically needed to complete the training, which can substantially shorten the overall training duration. Furthermore emerging contrastive learning frameworks that leverage self-supervised pre-training on large unlabeled molecular datasets to generate enriched and transferable molecular representations hold considerable promise for further improving predictive performance in data-scarce scenarios and may represent a powerful complement to conventional transfer learning approaches in future drug-target interaction studies (Dalkiran *et al*, 2026; Wang *et al*, 2022).

Our findings provide proof of concept evidence that Carbimazole, Crizotinib, and GSK-2018682 may have an inhibitory effect on mitochondrial 1C metabolism; however other on- and off-target effects cannot be ruled out. Further study will be required to determine how these drugs affect the enzymatic activity of MTHFD2 and SHMT2.

## Methods

We framed drug–target interaction (DTI) prediction as a ligand-based binary classification problem in which a deep neural network classifies each compound as active or inactive against a specific target protein. The general transfer learning framework, model architecture, and compound feature encoding used here follow our previously published approach (Dalkiran *et al*., 2023). Briefly, training consists of two stages illustrated in Figure 1A: in Stage I, a deep feedforward neural network (DFNN) is pre-trained on a large source dataset drawn from a related enzyme family; in Stage II, the pre-trained source model is fine-tuned on the small target-specific dataset using two modes (Figure 1B). Mode 1 (full fine-tuning) updates all network weights starting from the source model weights rather than random initialization. Mode 2 (feature transformer) freezes the early layers and updates only the output layer, using the source model as a fixed latent feature extractor. The best-performing architecture, FNN-2-Chemprop, uses 300-dimensional molecular representations learned by Chemprop (Heid *et al*, 2024). as input, with two hidden layers of 1,200 and 300 neurons, a learning rate of 0.0001, batch size of 256, and 100 training epochs. Full architecture details and hyperparameter selection are described in (Dalkiran *et al*., 2023) and at Supplementary GitHub page (https://github.com/MutluHamanakaLab/Transfer_Learning_Enabled_DTI_Prediction).

ChEMBL (V29) database was used to construct training and test datasets along with a data filtering method (Rifaioglu *et al*., 2020). The compounds were transformed into ECFP4 fingerprints. To reduce the model’s bias towards certain chemical families during training and tests, the compounds were clustered using RDkit’s clustering tool (Butina, 1999) with a Tanimoto coefficient cut-off value of 0.8. The datasets were balanced between active and inactive compounds, with duplicate entries consolidated and chemical diversity maximized through clustering technique. All datasets were created using the representation learning capabilities of Chemprop (data available at Supplementary GitHub).

### Fibroblast Culture

Normal human lung fibroblasts were purchased from Lonza (catalog number CC-2512) and maintained in Fibroblast Growth Medium 2 (PromoCell, C023020). Cells were plated at 1 × 10^5^ on 12 well plates for experiments. 24 hours after plating, cells were serum starved in DMEM (Gibco, 11054020) containing 0.1% bovine serum albumin (BSA), 5.5 mM glucose, 2 mM glutamine, and 1 mM pyruvate for 24 hours prior to treatment with 1 ng/mL TGF-β (Peprotech, 100-21 C). Predicted drugs (Figure 2) were added at the time of TGF-β treatment.

### Western Blot

Cells were lysed, and electrophoresis was performed as we previously described. Wells were lysed in 100 μL Urea Sample Buffer (8 M deionized urea, 1% SDS, 10% Glycerol, 60 mM Tris pH 6.8, 0.1% pyronin-Y, 5% β-mercaptoethanol). Lysates were run through a 28 gauge needle and were electrophoresed on Criterion gels (Bio-Rad) and transferred to nitrocellulose using a Trans-Blot Turbo (Bio-Rad) set to the Mixed MW program. Primary antibodies used were: Collagen 1 (Abcam, ab138492), Collagen 3 (Proteintech, 22734-1-AP), α-SMA (Sigma, A2547), GAPDH (Cell Signaling, 2118). Secondary antibodies were HRP-linked anti-rabbit and anti-mouse (Cell Signaling, 7074 and 7076, respectively). Blots were imaged using a ChemiDoc Touch (Bio-Rad).

### Gas Chroatography-Mass Spectrometry

HLFs grown for 48 hours in the presence or absence of TGF-β were washed in blood bank saline (Thermo, 23-293-184) and metabolites were extracted in 600 μL ice cold 80% Methanol (Fisher, A456 (MeOH), W6 (H_2_O)). The solution was vortexed and centrifuged at 21,000 × *g* for 20 minutes. 400 μL of each extract was transferred to a new tube and dried under nitrogen. Dried metabolites were derivatized in 16 μL Methoxamine reagent (Thermo, TS-45950) for 1 hr at 37 °C followed by 20 μL of 1% *N*-*tert*-butyldimethylsilyl-*N*-methyltrifluoroacetamide (Sigma, 394882) for 1 hr at 60 °C. Derivatized samples were analyzed with an 8890 gas chromatograph with an HP-5MS column (Agilent) coupled with a 5977B Mass Selective Detector mass spectrometer (Agilent). Helium was used as the carrier gas at a flow rate of 1.2 ml/min. One microliter of each sample was injected in split mode (1:8) at 280 °C. After injection, the GC oven was held at 100 °C for 1 min and increased to 300 °C at 3.5 °C/min. The oven was then ramped to 320 °C at 20 °C/min and held for 5 minutes. The MS system was operated under electron impact ionization at 70 eV and the MS source was operated at 230 °C and quadrupole at 150 °C. The detector was used in scanning mode, and the scanned ion range was 100–650 *m/z*. Peak ion chromatograms for metabolites of interest were extracted at their specific *m/z* with Mass Hunter Quantitative Analysis software (Agilent Technologies). Ions used for quantification of metabolite levels were as follows: Glycine *m/z* 246.

